# Species substitution in the meat value chain by high-resolution melt analysis of mitochondrial PCR products

**DOI:** 10.1101/2021.01.12.426171

**Authors:** Jane K. Njaramba, Lillian Wambua, Titus Mukiama, Nelson Onzere Amugune, Jandouwe Villinger

## Abstract

Food fraud in several value chains including meat, fish, and vegetables has gained global interest in recent years. In the meat value chain, substitution of high commercial-value meats with similar cheaper or undesirable species is a common form of food fraud that raises ethical, religious, and dietary concerns. The presence of undeclared species could also pose public health risks caused by allergic reactions and the transmission of food-borne or zoonotic pathogens. Measures to monitor meat substitution are being put in place in many developed countries. However, information about similar efforts in sub-Saharan Africa is sparse. In this study, we used PCR coupled with high-resolution melting (PCR-HRM) analysis targeting the three mitochondrial genes, cytochrome oxidase 1 (*CO1*), cytochrome b (*cyt b*), and *16S rRNA*, to detect species substitution in meat sold to consumers in Nairobi, Kenya’s capital city. Out of 107 meat samples from seven common livestock animals (cattle, goat, sheep, pig, chicken, rabbit, and camel), 11 (10.3%) had been substituted. Of 61 samples sold as beef, two were goat and one was camel. Of 30 samples sold as goat meat, four were mutton (sheep) and three were beef. One of nine samples purchased as pork was beef. Our results indicate that PCR-HRM analysis is a cost and time effective technique that can be employed to detect species substitution. The combined use of the three markers produced PCR-HRM profiles that successfully allowed the distinction of species. We demonstrate its utility not only in analysis of raw meat samples, but also of cooked, dried, and rotten samples, meat mixtures, and with the use of different DNA extraction protocols. We propose that this approach has broad applications in authentication of meat products and protection of consumers from food fraud in the meat industry in low- and middle-income countries such as Kenya, as well as in the developed world.

## 1. Introduction

Food fraud, the intentional act of adulterating food products, often for dishonest economic gain is an emerging concern in global trade as a crime against consumer rights, and due to the inherent risks posed to public health. Food fraud is largely perpetrated by counterfeit descriptions of products with respect to their weights, details of origin, types of processing, and constituents (ingredients) (Spink & Moyer, 2011). Food fraud has been reported in most value chains, including spices (Silvis et al., 2017), milk (Handford et al., 2016), edible oils (Yadav, 2018), cereals (Nasreen & Ahmed, 2014), vegetables (Panghal et al., 2018; Woolfe & Primrose, 2004), and meat (Chuah et al., 2016; Ren et al., 2017), whereby fraudulent substitution of ingredients or adulteration of products with similar but cheaper options has been highlighted as major malpractice.

In the meat industry, the major fraudulent practice entails substitution of meats of high commercial value with those from cheaper or undesirable species (Chuah et al., 2016; Farag et al., 2015). Major global incidences of species substitution have been reported, such as the horsemeat scandal in the UK and Ireland where beef was substituted with horse meat (Di Pinto et al., 2015) and in China, where mutton was substituted with murine meat (Fang & Zhang, 2016). In Kenya, substitution of beef and chevron with bushmeat (Kimwele et al., 2012; Ouso et al., 2020) in addition to reports of species-substitution (Kenya Markets Trust, 2019) necessitates further study of efficient methods of detecting this malpractice in meat value chains. Species substitution in meat products inhibits fair trade (Ballin et al., 2009) and raises ethical and religious concerns where species substitutes sold are considered offensive (Al-Kahtani et al., 2017; Chuah et al., 2016). Undeclared meat species are also a health liability to those with allergies (Di Pinto et al., 2015) and are associated with public health safety risks such as those posed by foodborne or zoonotic diseases. Substituting species utilized are frequently acquired from unconventional sources, such a wildlife (bushmeat), and could have been subjected to unhygienic handling and may not have undergone quality checks like meat inspection (Alarcon et al., 2017a).

Detection of adulteration in the meat value chain relies on analytical techniques such as chromatography, mass spectrophotometry, imaging, and serology to identify particular contaminants, proteins, metabolites, and validate authenticity (Abbas et al., 2018; Cawthorn et al., 2013; Farag et al., 2015). However, for analysis of species substitution, DNA-based techniques have been increasingly adopted due to the inherent limitations in specificity and sensitivity associated with the aforementioned techniques (Abbas et al., 2018), leading to the recognition of “Food Forensics” as a tool to investigate food fraud (Woolfe & Primrose, 2004). The use of DNA to identify species on the basis of universal barcoding markers has been reliably tested (Farag et al., 2015; Ouso et al., 2020). These methods have evolved from the more conventional PCR-based techniques (Farag et al., 2015; Sakaridis et al., 2013) to more novel techniques including PCR coupled with high-resolution melting (HRM) analysis. PCR-HRM allows for discrimination of DNA variants by detection of nucleotide sequence differences such as single nucleotide polymorphisms (SNPs) and insertions and deletions (indels) based on their melting profiles, hence enabling genotyping of species (Reed et al., 2007). PCR-HRM analysis has also been used to identify vertebrate species in insect blood-meals (Omondi et al., 2015), bushmeat (Ouso et al., 2020), and adulteration of buffalo meat (Sakaridis et al., 2013). Nevertheless, the accuracy of results for DNA-based analyses depends fundamentally on obtaining quality DNA to identify the species origins of frozen, cooked, processed, rotting, or mixed meat products (Cawthorn et al., 2013; Farag et al., 2015; Sakaridis et al., 2013).

Studies have demonstrated the potential application of DNA sequencing to identify adulteration in meat products based on *CO1* and *cyt b* genes (Kimwele et al., 2012; Song’oro et al., 2012; Mbugua et al., 2014; Bourguiba-Hachemi & Fathallah, 2016). While useful, the need for elaborate and relatively expensive post-PCR procedures, such as DNA sequencing, severely limits their usefulness in routine monitoring of meat fraud in Kenya and other low-resource settings. Additionally, the effect of different physicochemical states of meat on PCR efficiency remains understudied. Therefore, we studied the utility of PCR-HRM analysis targeting *CO1*, c*yt b*, and *16S rRNA* genes to investigate species substitution in Nairobi, Kenya. We also aimed to test the effect of various meat matrices (e.g. fresh, dried, cooked, or rotten meat), different DNA extraction protocols, and mixed-meat samples on species identification by PCR-HRM analysis.

## 2. Materials and Methods

### 2.1 Meat samples

We purchased 107 meat samples in November 2018 from randomly selected stalls in Nairobi’s major meat wholesale market (Burma market), and butcheries in the surrounding estates of Eastleigh, Kariokor, Kaloleni, Mukuru Village, Mathare, Jerusalem, Jericho, Ngara, and Makongeni (Supplementary Figure 1). Meat samples of species commonly bought by households were purchased, including 61 cattle, 30 goat, three camel, nine pig, and four chicken samples. Each 250-g sample was packed separately and transported in cooler boxes with ice packs to the lab. Sub-samples (1 g) were carefully excised from the internal portion of each sample to obtain two replicates. Sterile blades and fresh gloves were used for each sample on a sterile surface. The replicates were then stored in 2-ml cryovials at −80°C pending DNA extraction. Twenty-four reference meat samples of known vertebrates archived from a previous study (Ouso et al., 2020) were used as positive controls (Supplementary Table 1). Genomic DNA was DNA extracted from the sub-samples using the ISOLATE II Genomic DNA Extraction Kit (Bioline, UK) following the manufacturer’s instructions.

### 2.2 Identification of vertebrate sources of meat by PCR-HRM

To identify the vertebrate species, DNA extracts from the test samples and positive controls (reference samples) were then analyzed by PCR-HRM of vertebrate mitochondrial *cyt b*, *CO1*, and *16S rDNA* as previously described (Ogola et al., 2017; Omondi et al., 2015; Ouso et al., 2020). Briefly, 10-μl PCR reactions were set up, each comprised of 1X HOT FIREPol^®^ EvaGreen^®^ HRM Mix no ROX (Solis BioDyne, Tartu, Estonia), 0.5 μM of both forward and reverse primers, 20 ng DNA template and nuclease free water. Each run included a negative control where ddH2O was added in place of DNA template. The PCR-HRM analyses were carried out in a RotorGene Q thermocycler (Qiagen, Germany) as described by Ouso *et al.* (2020). Briefly, the cycling conditions involved an initial hold at 95°C for 15 minutes, followed by 40-45 cycles of denaturation at 95°C for 20 seconds, annealing for 20 seconds at 56°C, and an extension step at 72°C for 30 seconds. This was followed by the final extension step, an additional 5 minutes at 72°C. The amplicons were then gradually melted from 75°C to 95°C while recording fluorescence at after two seconds at 0.1°C increments. Melt rate and normalized HRM graphs were generated from the fluorescence data, using the Rotor-Gene Q Series Software (2.3.1 build 49). Meat-source species were distinguished by analyzing the melt rate (melting temperature (Tm) peaks) and normalized profiles of the test samples against those of the reference species. For species with single Tm-peaks, we also examined the Tm-deviation from the control, with similar species expected to have Tm shifts of < 1°C.

### 2.3 Analysis of various physicochemical treatments of meat on PCR-HRM

To study the effect of various physicochemical conditions of meat samples on species identification by PCR-HRM analysis, we utilized sub-samples from our collection of positive controls (see Supplementary Table 1). Sub-samples from each of goat (*Capra hircus*), sheep (*Ovis aries*), pig (S*us scrofa domesticus*), and chicken (*Gallus gallus domesticus*), cattle (*Bos taurus*) and camel (*Camelus dromedarius*) were exposed to different treatments to simulate fresh, dried, cooked (microwaved), and rotting/decomposed meat. This was achieved by obtaining four replicates weighing 60 mg from each sub-sample and treating them as follows: the first replicate was used as the fresh meat with no treatment was applied, the second replicate was dried in an oven at 65°C for 2 hours, the third replicate was heated in a microwave oven for 12 minutes to simulate cooking, and the fourth replicate was left on the lab bench for 72 hours to decompose. Genomic DNA was extracted from the replicates of all samples as described above in section 2.1, followed by PCR-HRM of the *CO1* gene, *cyt b*, and *16S rRNA* genes as described in section 2.2.

### 2.4 Analysis of effect of different extraction protocols

To study the impact of different DNA extraction protocols on species identification by PCR-HRM, sub-samples were obtained from two cattle, four goats, one sheep and two camels as described in section 2.3 above and subjected to four extraction protocols. Each sub-sample was divided into four 50-mg replicates and DNA was extracted as follows: The first replicate was extracted using the ISOLATE II Genomic DNA Kit as described in 2.1 and the second using the DNeasy Blood and Tissue Kit protocol (Qiagen, Valencia, CA) according to the manufacturer’s guidelines. The third replicate was extracted using a lab-optimized protocol described by Kipanga and co-workers (Kipanga et al., 2014). The fourth replicate was extracted using a modified version of the aforementioned protocol, where proteinase K was omitted during the cell-lysis step. The extracted DNA was then standardized to 10 ng/μl and analyzed using PCR-HRM of *CO1*, *cyt b*, and *16S rRNA* as described in section 2.2. The melt profiles were then compared to check for any differences in melt temperature or profile due to extraction protocol differences.

### 2.5 Analysis of species admixtures in meat by PCR-HRM

We investigated whether PCR-HRM could be successfully used to identify mixed species in meat, which is a common adulteration in processed meat. The following mixtures were prepared from the reference samples: cattle + sheep; sheep + goat; cattle + goat; cattle + camel; chicken + pork; and chicken + Nile perch. In each case, triplicates containing 50 mg of each of the two species in the combinations above were placed into separate tubes. Genomic DNA was then extracted from the individual triplicates using the ISOLATE II Genomic DNA Kit as described in section 2.1. This was followed by PCR-HRM analysis of the three mitochondrial markers (*CO1*, *cyt b*, and *16S rRNA*) as previously described. DNA extracts from individual reference samples, i.e. individual positive controls of cattle, sheep, goat, chicken, pork, camel, and Nile perch, were also analyzed alongside the mixed samples.

### 2.6 Vertebrate species confirmation and statistical analysis

To confirm the vertebrate species in the meat samples, the DNA was amplified using primers that target a longer segment (750 bp), the barcoding region of the *CO1* gene, as described previously (Ivanova et al., 2012; Ouso et al., 2020). This involved amplification using conventional PCR using 15-μl reaction volumes which included 1X HOT FIREPol® Blend Master Mix (Solis BioDyne, Tartu, Estonia), 0.5 μM concentrations of both forward (5’-TCT CAA CCA ACC ACA ARG AYA TYG G-3’) and reverse (5’-TAG ACT TCT GGG TGG CCR AAR AAY CA-3’) primers and 2 μl of DNA template. The cycling conditions were those described by Ouso *et al.* (Ouso et al., 2020). The resulting amplicons were cleaned using the ExoSAP-IT protocol (USB Corporation, Cleveland, OH) and sequenced at Macrogen Inc. (Netherlands). Sequences were analyzed using Geneious version 11.1.5 (Kearse et al., 2012; Lee et al., 2012) and queried against the GenBank nr database (http://www.ncbi.nlm.nih.gov/) using the Basic Local Alignment Search Tool (BLAST; Altschul et al., 1990) and the Barcode of Life Database (BOLD; http://www.boldsystems.org; Ratnasingham & Hebert, 2007). The statistical software NCSS 2020 (NCSS, Kaysville, Utah, USA; https://www.ncss.com/) was used to create box plots of the variance in melting temperatures observed using different extraction conditions or physicochemical treatments.

## 3. Results

### 3.1 Vertebrate sources of meat sold in butcheries in Nairobi

PCR-HRM analysis of the *cyt b, CO1*, and *16S rRNA* genes (Supplementary Figure 2) of 107 meat samples revealed the vertebrate sources as 62 cattle (57.94 %), 25 goats (23.36%), eight pigs (7.47%), four camels (3.74%), four chicken (3.74%), and four sheep (3.74%). Identifications were confirmed by sequencing of the long *CO1* barcode amplicons. Eleven (10.3%) meat samples were misidentified by sellers. Of 61 samples sold as beef, two were substituted with goat meat and one with camel meat. Of 30 samples sold as goat meat, four were mutton (sheep meat) and three were beef. One of the nine samples purchased as pork was beef (Figure 1). Pair-wise comparison of the amplicons allowed for the distinction of different species using the three primers (Figure 2).

**Figure 1:**
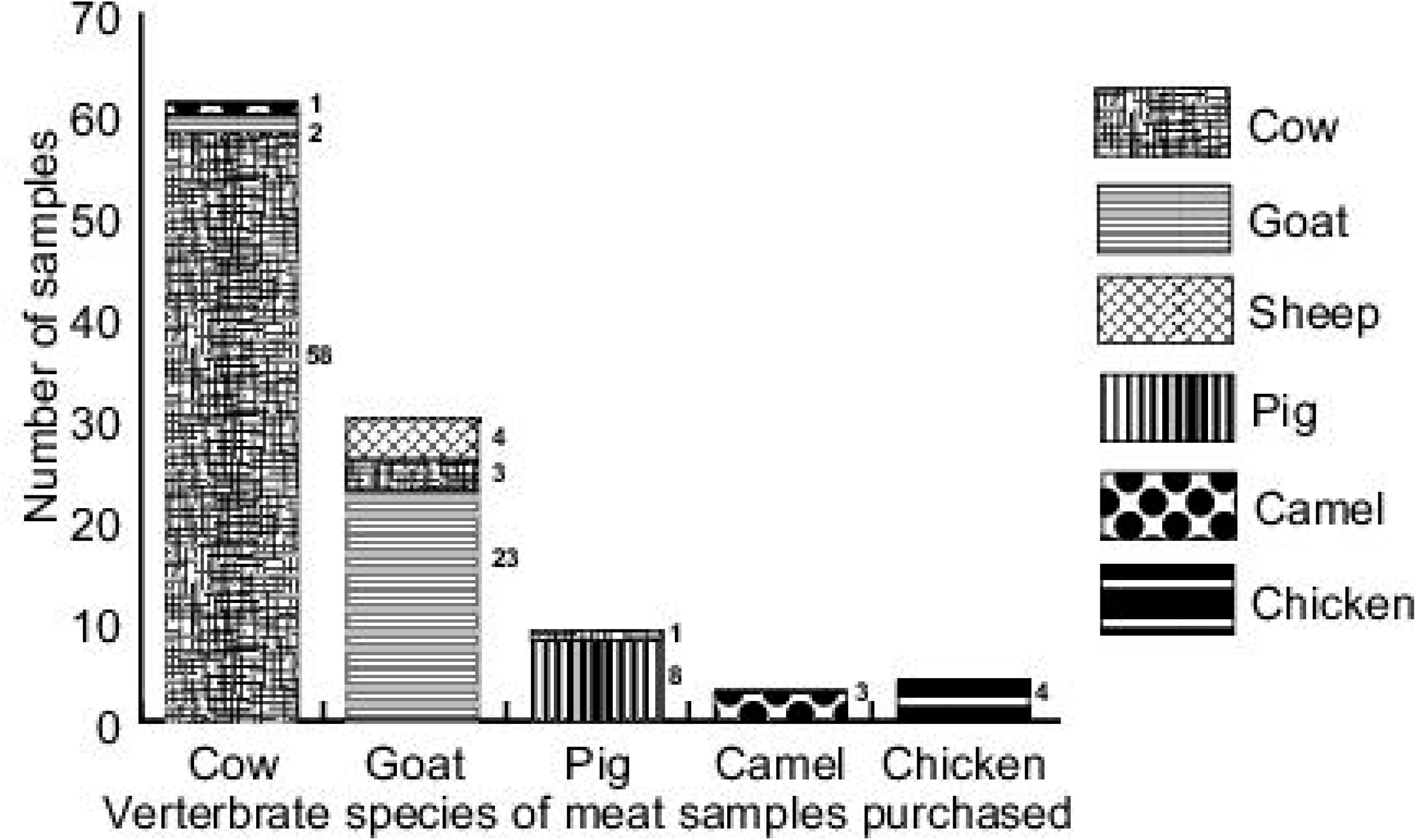
Species substitution of meat sampled in Nairobi. Stacked bar graph showing vertebrate species of meat identified by PCR-HRM against the identity of the species purchased in the meat market. Numbers against each species refers to *n*, the number of species identified using PCR-HRM. Species substitution was identified in cattle, goat and pig samples.

**Figure 2:**
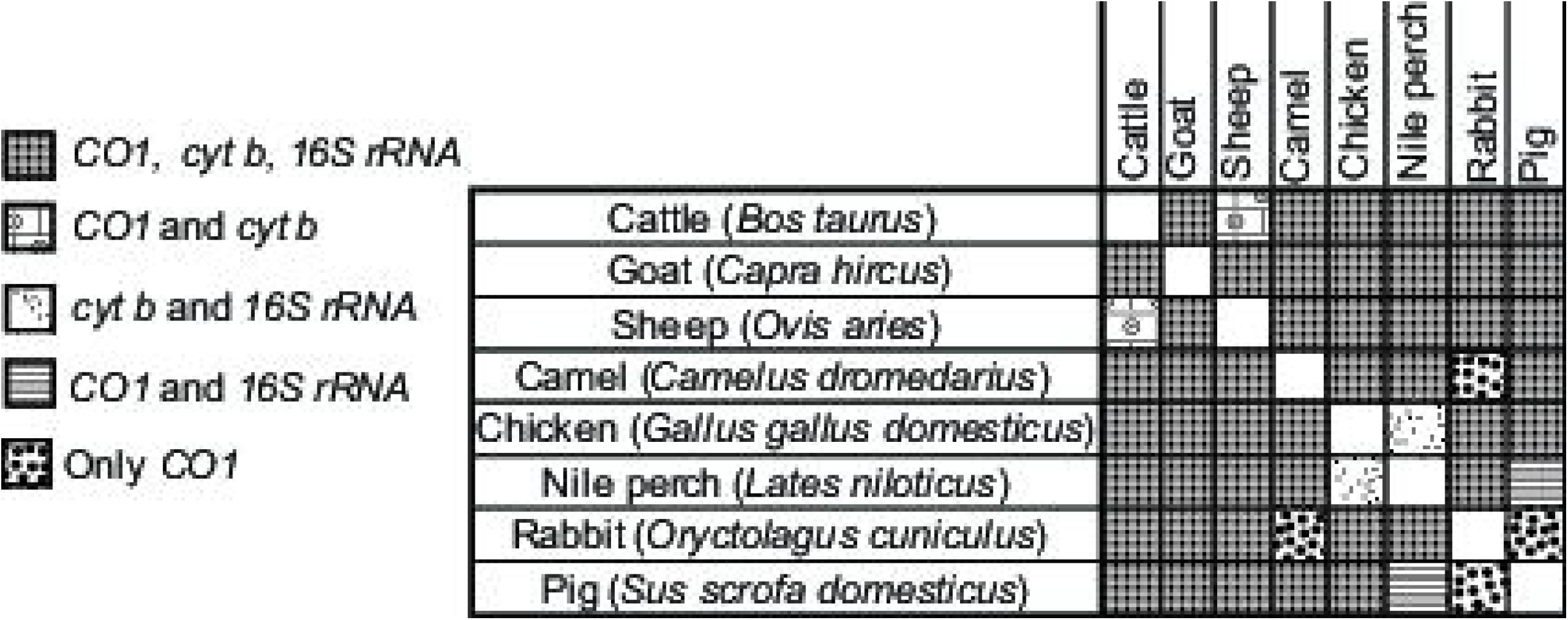
Pairwise discrimination of vertebrate sources of meat by PCR-HRM analysis: Three mitochondrial markers *CO1*, *cyt b* and *16S rRNA* were compared. The ability of these markers to distinguish eight different vertebrate species commonly consumed in Kenyan households was compared and the summary matrix was generated.

### 3.2 Effect of physicochemical condition of meat samples on vertebrate species identification by PCR-HRM

We found that the application of various treatments had minimal effect on the HRM melt profiles of the respective PCR amplicons. All samples irrespective of the physicochemical condition, were amplified by at least one of the markers and the vertebrate species could be reliably identified despite slight shifts in the melting temperature (Tm) of the resulting amplicons. Amplification was highest in raw samples, with all of them being successfully detected using all three markers. However, we observed relative reduction in the amplification of the markers different treatments. In the *CO1* gene, 8/16 of the microwaved, 2/16 of the rotten, and 4/16 of the oven-dried samples did not amplify, while in the assay targeting the *cyt b* marker, 10/16 of the microwaved samples, 2/16 of the oven-dried samples, and 1/16 rotten samples did not amplify. The *16S rRNA* did not amplify for 6/16 of the microwaved and 1/16 of the oven-dried samples.

Comparing the melting temperature Tm of PCR amplicons obtained from oven-dried, cooked, and rotten meat with the raw samples indicated slight shifts in the Tm. The *cyt b* marker had the highest impact on the Tm as observed by the range in Tm shift, when samples exposed to the different conditions, whereas the Tm of the *CO1* marker was least affected by meat treatment. The shift in Tm from the raw-meat controls was < 1°C in all markers for all samples except one microwaved cattle sample, which had a *16S rRNA* Tm shift of +1.63°C. Consequently, the widest range in primary-peak Tm was seen in microwaved samples, followed by those degraded, and oven-dried showing the least variation from the Tm observed in the control (raw) meat samples (Figures 3 and 4). We noted that the amplification of the *16S rRNA* marker resulted in single peaks in all the species tested, whereas the *cyt b* and *CO1* markers resulted in prominent secondary peaks that help to distinguish cattle, sheep, and chicken (Supplementary Figure 3). Camel samples had double peaks only in the *cyt b* region, whereas pig only had multiple peaks in the *CO1* region.

**Figure 3:**
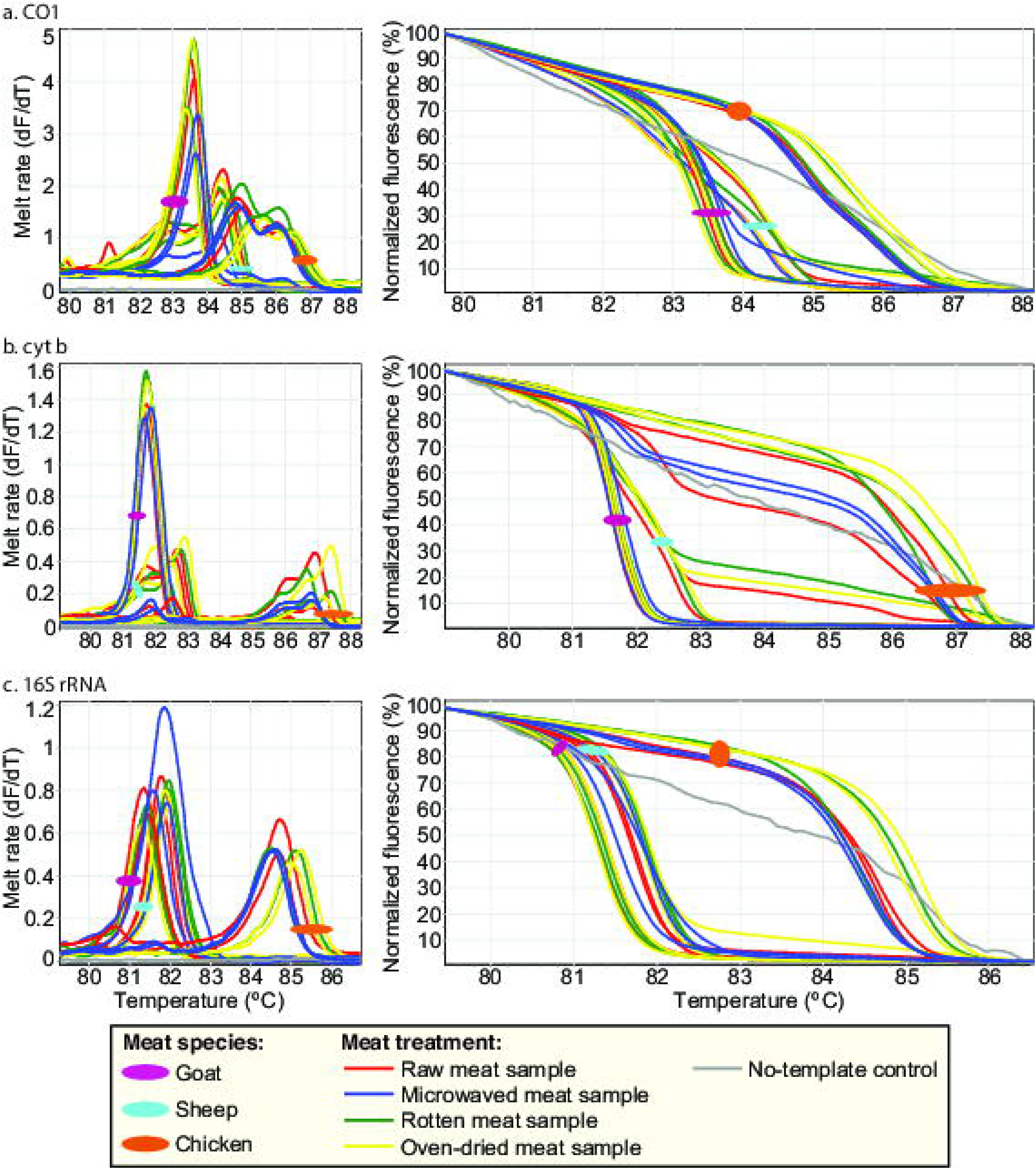
PCR-HRM profiles of representative reference samples exposed to different physicochemical conditions. Goat, sheep and chicken meat samples were exposed in replicate to different conditions; raw, rotten, oven-dried and microwaved. Their PCR-HRM profiles were then assessed using *CO1*, *cyt b* and *16S rRNA*. For each marker, the HRM profiles are represented as melt rates and normalized HRM profiles. Melt rates are represented as change in fluorescence units with increasing temperatures (dF/dT) and HRM profiles are represented as percent fluorescence with increase in temperature for a) *CO1*, b) *cyt b*, and c) *16S rRNA* markers.

**Figure 4:**
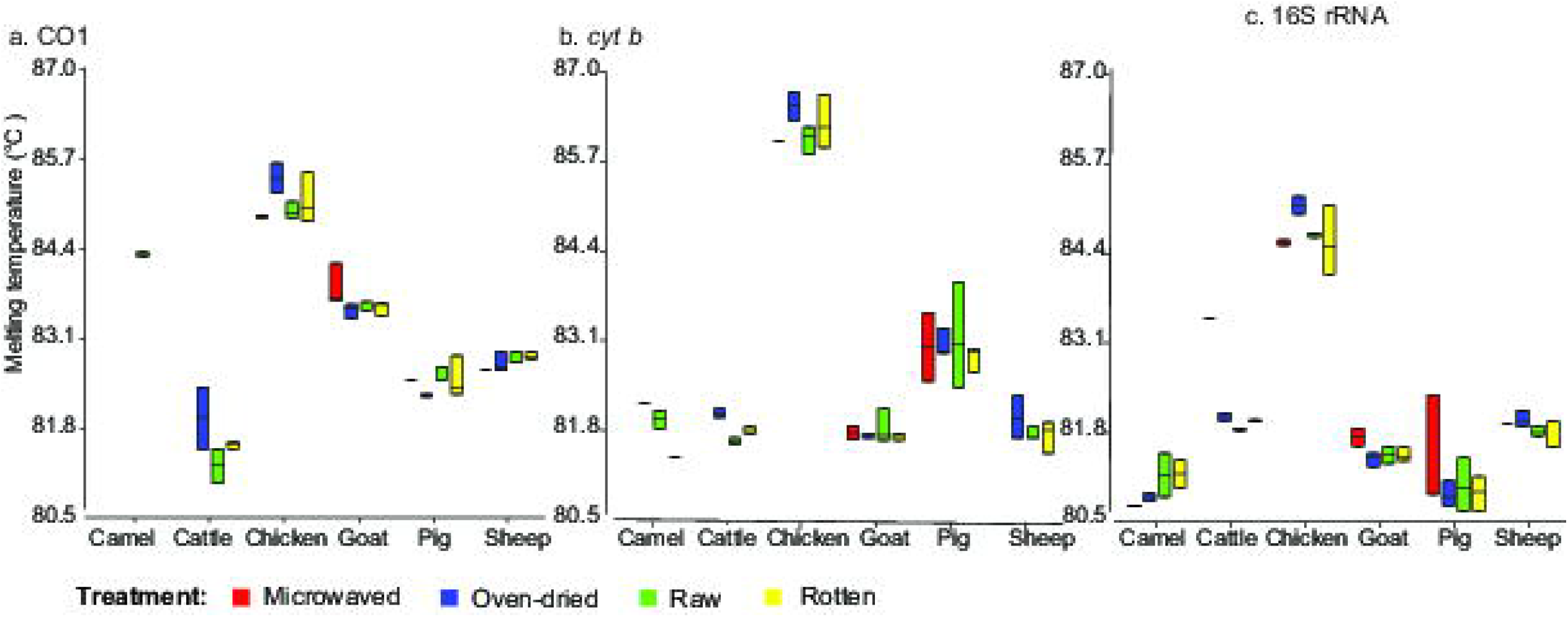
Box plots of peak melting temperatures (°C) of meat samples exposed to different physicochemical conditions. Replicate meat samples from goat, sheep, pig, cattle, camel, and chicken were exposed to different conditions; raw, rotten, oven-dried, and microwaved. The peak PCR-HRM melt rate temperatures were plotted for a) *CO1*, b) *cyt b*, and c) *16S rRNA* markers.

### 3.3 Effect of different DNA extraction protocols on PCR-HRM

Using the different DNA extraction protocols, we observed similar melt profiles with minimal Tm shifts (< 1°C). The *CO1* marker had the widest range in Tm, followed by *cyt b* (Figure 5; Figure 6). The use of different extraction protocols did not result in overlapping of profiles of any of the species analyzed. All markers could be used to distinguish the species of the samples, regardless of extraction protocol used, with the exception of cattle and sheep which yielded similar HRM melt profiles with *16S rRNA* marker (Supplementary Figure 4).

**Figure 5:**
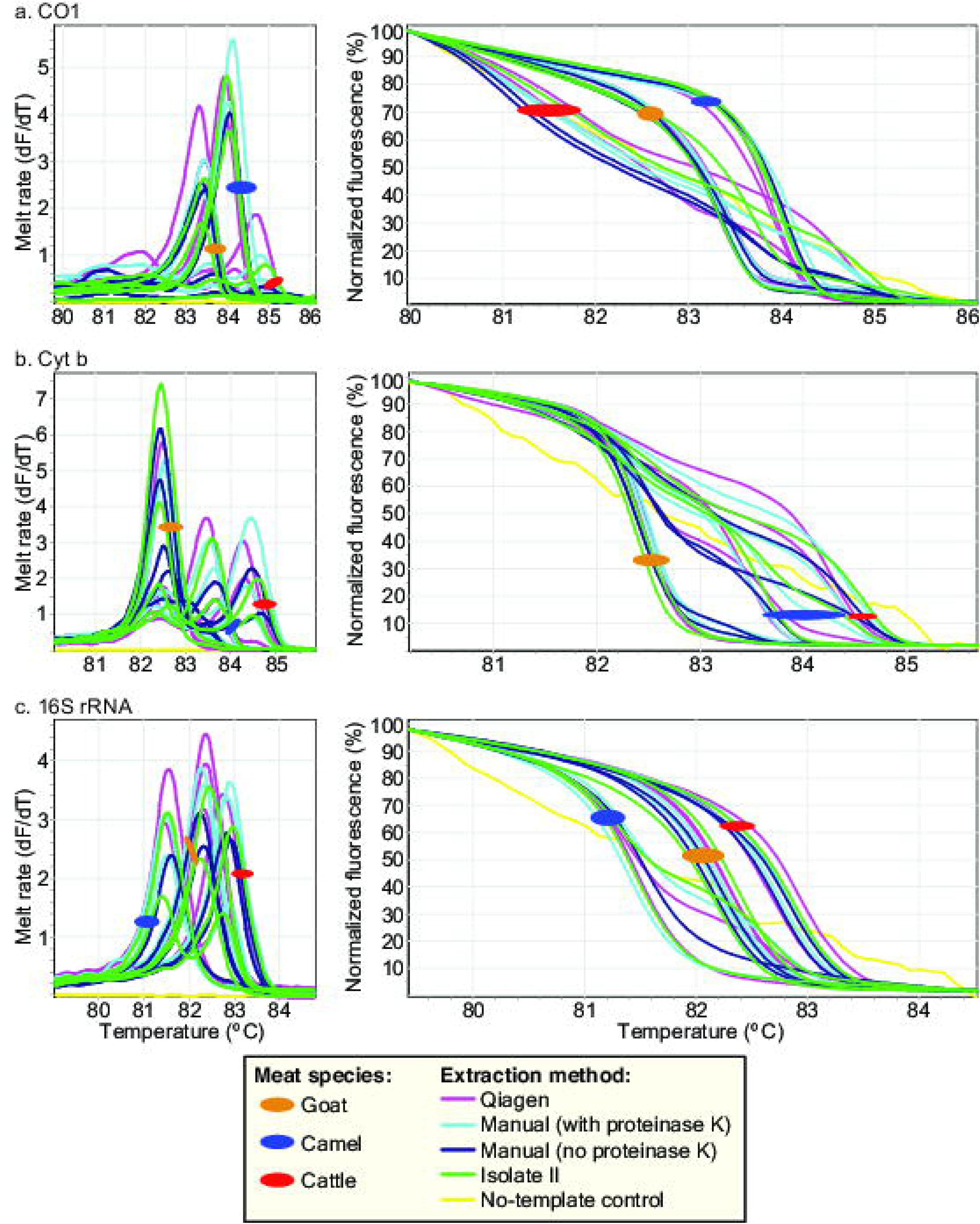
Effect of different DNA extraction protocols on PCR-HRM profiles of selected reference vertebrate species. DNA were extracted from goat, camel and cattle meat samples using four different extraction protocols in replicates. These protocols included two kits: DNeasy Blood and Tissue Kit protocol, the ISOLATE II Genomic DNA Extraction Kit and two manual extraction protocols. Their PCR-HRM profiles were then assessed using *CO1*, *cyt b* and *16S rRNA*. For each marker, the HRM profiles are represented as melt rates and normalized HRM profiles. Melt rates are represented as change in fluorescence units with increasing temperatures (dF/dT) and HRM profiles are represented as percent fluorescence with increase in temperature for a) *CO1*, b) *cyt b*, and c) *16S rRNA* markers.

**Figure 6:**
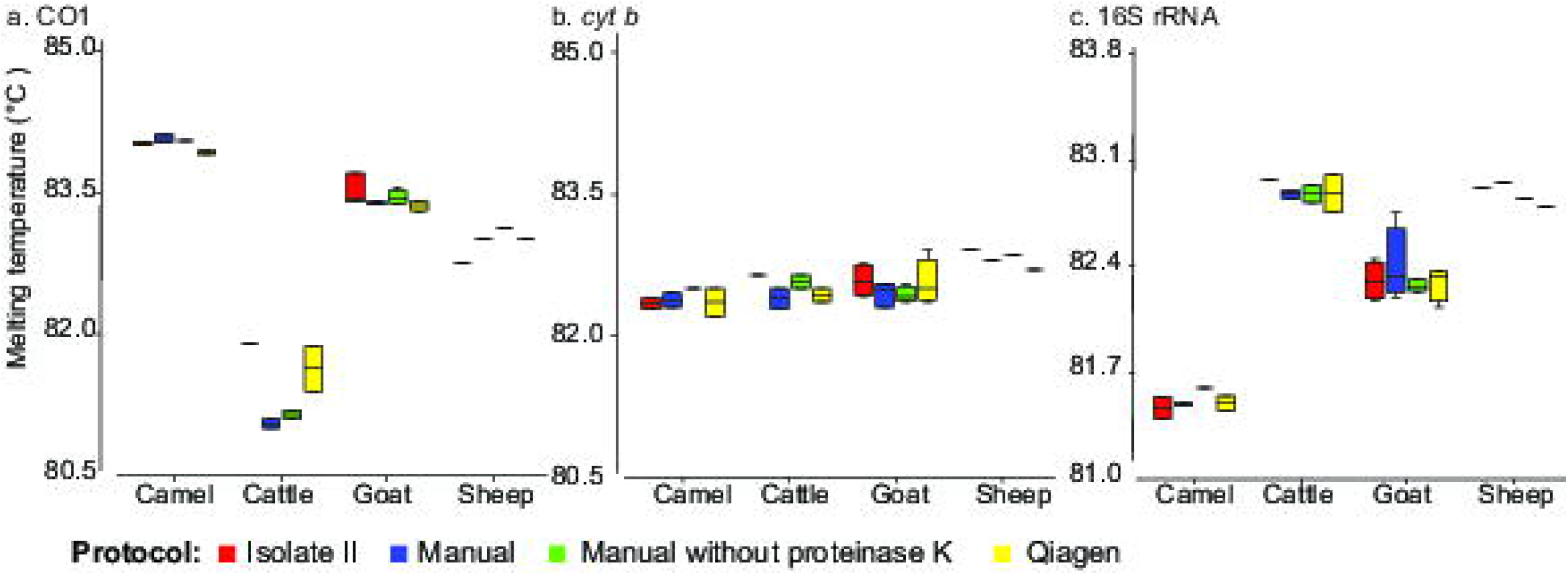
Box plots of peak melting temperature (°C) seen in meat samples extracted using different extraction protocols. DNA from four vertebrate species (two cattle, four goats, one sheep and two camels) were extracted in replicate using two commercial kits; DNeasy Blood and Tissue Kit protocol, the ISOLATE II Genomic DNA Extraction Kit, and two manual extraction protocols. The peak PCR-HRM melt rate temperatures were plotted for a) *CO1*, b) *cyt b*, and c) *16S rRNA* markers.

### 3.4 Distinction of species in mixed meat samples using PCR-HRM

Amplification targeting the marker *16S rRNA* gave the best resolution in distinguishing the individual vertebrate species in mixed meat samples (meat samples with 2 or more vertebrate species). The only mixed samples that could not be determined using the *16S rRNA* marker were mixtures of cattle and sheep meat, which could however be distinguished by the *cyt b* marker (Figures 2 and 7). The *cyt b* marker, clearly resolved white meat mixtures including chicken and pork and Nile perch and pork, with individual HRM curves corresponding with the composite vertebrate species. However, differentiating sources of red meat using the *cyt b* marker was limited. For instance, all mixtures that contained goat meat only showed the melt profile of goat *cyt b* profile. The melting profiles obtained from the *CO1* marker showed slight variations between pure samples (meat samples with one vertebrate species) and the mixed meat samples.

**Figure 7:**
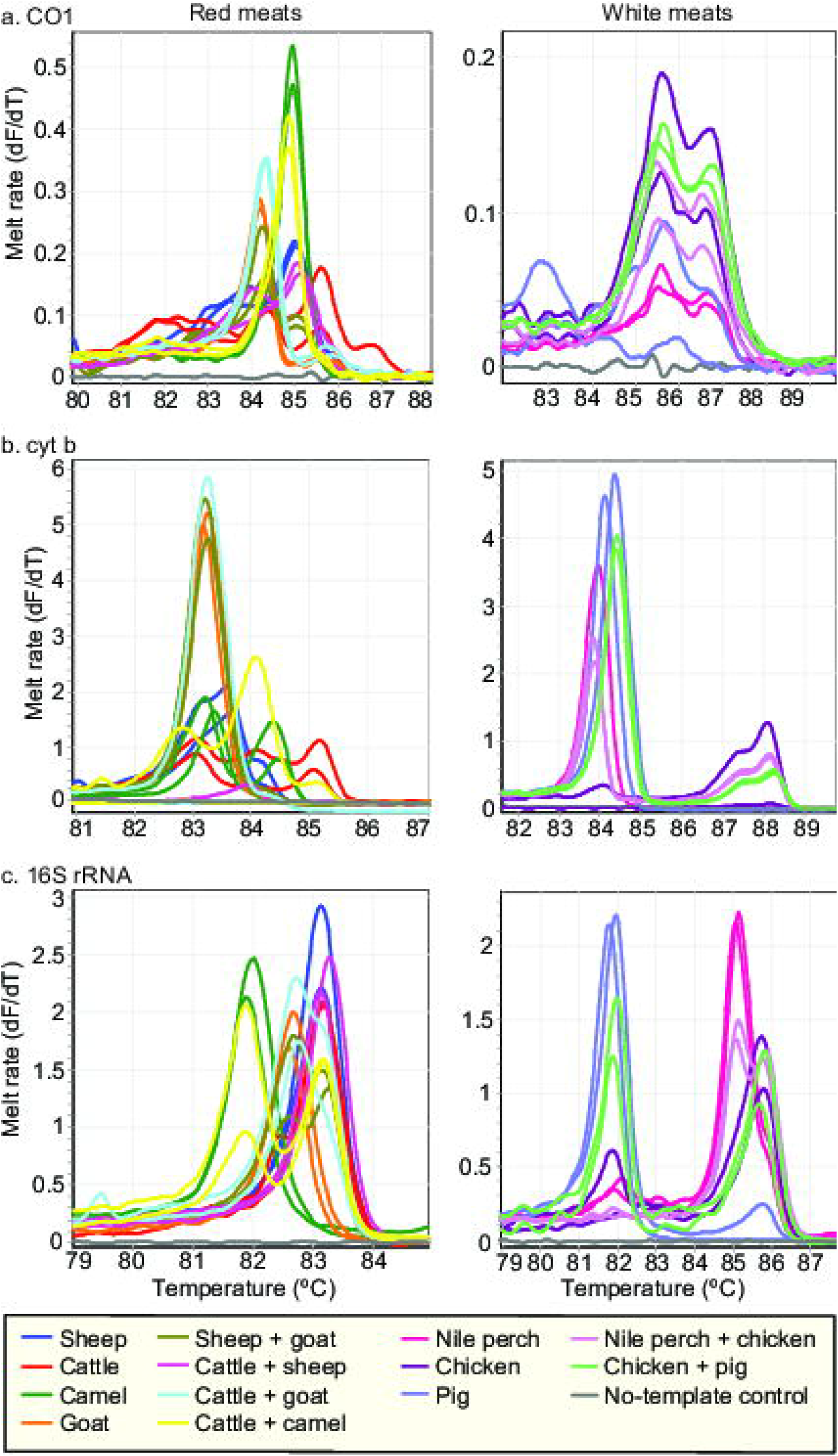
PCR-HRM melt rate profiles of pure (single species) and mixed meat samples assessed using three mitochondrial markers *CO1*, *cyt b* and *16S rRNA*. The left column represents red meat sources (sheep, goat, cattle, and camel) and their corresponding mixtures, whereas the right column represents DNA from white meat sources (Nile perch, chicken, and pig) and their mixtures. Distinct melt rates are represented as change in fluorescence units with increasing temperatures (dF/dT) for a) *CO1*, b) *cyt b*, and c) *16S rRNA* markers.

## 4. Discussion

This study revealed significant levels of species substitution in products retailed in Nairobi’s major meat market. Goat meat had the highest levels of substitution, with mutton and beef being used as alternatives. Our results compare with those of Cawthorn and co-workers (2013), who reported detection of mutton and beef as common substitutes in the meat value chain in South Africa. In Kenya, this substitution is likely to be driven by the relatively higher price of goat meat (USD 5.5 – 6.0 per kg) relative to mutton and beef (USD 3 – 4.5 per kg) (Alarcon et al., 2017b). Goat meat is preferred for preparing “*Nyama Choma*”, a roasted meat delicacy in eastern Africa that is increasingly being consumed in high quantities (Gorski et al., 2016). Growing evidence associating consumption of beef and mutton with a severe allergic reaction termed “*midnight anaphylaxis*” implies that substitution with these meats from these species may pose a health risk to susceptible populations (Gray et al., 2016). While the detection of these undeclared species could be a result of unintentional cross-contamination, such as from dirty knives or surfaces, we accounted for these potential mishaps by testing inner parts excised from the raw meat samples. These results highlight the need for intensified surveillance of species substitution in the meat value chain in Kenya. Targeted surveillance may also be applied to goat value chains and other high value species where adulteration could be linked to fraudulent financial gain.

This study demonstrates the utility of PCR-HRM for detecting multiple vertebrate species in meat products. We were able to detect composite vertebrate species in mixed meat samples by using different combinations of *CO1*, *cyt b* and *16 rRNA.* Adulteration of meat with products from multiple species is increasingly being reported (Ali et al., 2015; Di Pinto et al., 2015; Izadpanah et al., 2018; Kitpipit et al., 2014; O’Mahony, 2013), raising the demand for affordable and faster techniques for their detection. Many studies describing multi-species analysis of vertebrates in meat have utilized multiplex PCR (Ali et al., 2015; Balakrishna et al., 2019; Izadpanah et al., 2018; Kitpipit et al., 2014; Li et al., 2019; Liu et al., 2019; Qin et al., 2019; Wang et al., 2020). While useful, multiplex PCR requires use of expensive probes and post-PCR procedures such as agarose gel electrophoresis for size separation of amplicons and/or DNA sequencing, thereby increasing analysis time, cost, and risk of cross-contamination. The PCR-HRM technique is hinged on measuring the rate of dissociation of PCR amplicons from double-stranded to single-stranded forms when subjected to gradual heating. Therefore, it allows real-time detection while minimizing downstream steps and costs. In a previous study, we demonstrated the utility of PCR-HRM in distinguishing up to 32 vertebrate species (Ouso et al., 2020), whereby DNA sequencing was performed only on representative samples for purposes of species confirmation, or to investigate samples with questionable or novel melt profiles.

This study shows that PCR-HRM would be particularly useful in investigating admixture of vertebrate species in commercial processed meat products such as sausages, kebabs, meatballs and hams, which are frequently adulterated with multiple undeclared meats during processing (Wang et al., 2020). In performing such analysis, our findings underpin the need to employ more than one marker to ensure accurate species identification in meat admixtures. Over-reliance on a single marker could result in overlaps in melt curves of different species resulting in decreased sensitivity and poor resolution of meat mixtures (Lopez-Oceja et al., 2017). Due to the unpredictability associated with meat mixtures, we recommend the use of multiple mitochondrial markers, coupled with careful design and selection of primers for PCR-HRM studies.

The application of various treatments to the meat samples allowed us to mimic the conditions and states of degradation that meat samples may be found in due to post-slaughter changes from cooking, sun-drying, or rotting (Ramanan & Khapugin, 2017). This was important because the efficiency of PCR is sensitive to inhibitors that may potentially affect amplification and amplicon melt temperatures (Tm). The melting profiles were similar across raw, cooked, and rotten meat samples with most samples having Tm shifts of < 1°C. However, exposure to heat treatment in the microwaved samples resulted in lower amplification rates and increased the *16S rRNA* marker Tm of one cattle sample by > 1°C. A shift of this magnitude was however, not evident when targeting the *CO1* marker showing that the use of this marker in conjunction with the others will improve species identification in meats exposed to heat post-slaughter. Our findings also imply that although reliable species distinction requires a Tm shift of < 1°C, it can be more in treated samples, hence the need for further studies to optimize acceptable thresholds. Nonetheless, across markers, species identification by PCR-HRM is reproducible despite varying physicochemical states of the meat samples.

Our results also demonstrate that the use of different DNA extraction protocols yielded only slight variations between same-species samples. Notably, the deviation from the control DNA isolation kit (the ISOLATE II commercial kit) was minimal (< 1°C), with the lab-optimized protocol having a slightly wider range in melting temperature compared to the other protocols used. The variation in melting temperature across different protocols was likely caused by the difference in salt concentrations of the DNA yielded. Cations such as Mg^2+^ and Na^+^ interact with the highly charged DNA polyanion, favoring DNA-melting in conditions with lower Na^+^ concentrations (Ouso et al., 2020; Tan & Chen, 2006). Nevertheless, despite the marginal amplicon Tm shifts, melting profiles did not vary, confirming that PCR-HRM can be used with various DNA extraction protocols.

While PCR-HRM allows for efficient and reliable differentiation of species substitution in the meat value chain, it is not without limitations. PCR-HRM relies on identification of vertebrate species against reference samples that are run alongside the assays as positive controls. However, the samples selected as references may be subjective, determined by what researchers may deem as important, hence leaving other species in the initial analysis. Nevertheless, our previous studies show that PCR-HRM analysis provides a degree discovery of novel/uncharacterized species, which can be identified by DNA sequencing (Ouso et al., 2020). Furthermore, the mitochondrial markers *CO1* and *cyt b*, are often used singly in species identification (Di Pinto et al., 2015; Lopez-Oceja et al., 2017; Izadpanah et al., 2018), but our results indicate the need to use multiple markers in tandem for accurate species identification. Finally, mitochondrial markers which are commonly used for DNA barcoding of species, are not appropriate for quantification of species adulteration in meat mixtures because there are major differences in gene copies of mtDNA markers in different species (Ballin et al., 2009; Cai et al., 2017).

## 5. Conclusions

This study demonstrates the utility of PCR-HRM for efficient and reliable detection of species substitution as a form of fraud in the meat value chain. We utilized this technique to identify species substitution in Nairobi’s major meat market. We also showed that PCR-HRM is a robust technique, providing reproducible results in both single or mixed species samples despite variations in physicochemical properties or presence of multiple DNA extraction protocols. Our study shows that PCR-HRM enables identification of vertebrate species in meat samples without having to perform extensive DNA sequencing on most of the samples, making it the molecular tool of choice for surveillance of food fraud in low-resource settings such as those found in sub-Saharan Africa and other developing economies. Finally, this work demonstrates the importance of using the multiple mitochondrial markers including *CO1*, *cyt b*, and *16S rRNA* to accurately distinguish species.

## Acknowledgements

We acknowledge the technical support of Nancy Kagendi, Edward Makhulu and Joseph Oundo of *icipe’s* ML-EID laboratory, and James Kabii of Molecular Biology and Bioinformatics Unit. We are thankful to Daniel Ouso for the provision of vertebrate positive controls used in the study. We also acknowledge the logistic support of Faith Kyengo, and Margaret Ochanda of the Capacity Building Unit at *icipe*, and Daudi Mutua of the Procurement Unit. We also thank Vivian Kerubo, of the Geo-Information Unit, *icipe* for her assistance in making the map.

## Author Contributions

**Jane Njaramba**: Investigation, Writing original draft, Data curation, Formal analysis. **Lillian Wambua**: Conceptualization, Funding acquisition, Writing original draft, Methodology, Formal analysis, Supervision. **Titus Mukiama**: Supervision. **Nelson Amugune**: Supervision. **Jandouwe Villinger**: Funding acquisition, Formal analysis, Methodology, Supervision, Visualization. **All authors:** Writing-Reviewing and Editing.

